# Cargo and biological properties of extracellular vesicles released from human adenovirus type 4-infected lung epithelial cells

**DOI:** 10.1101/2025.08.11.669368

**Authors:** Alessio Noghero, Stephanie Byrum, Chioma Okeoma, Adriana E. Kajon

## Abstract

Extracellular vesicles (EVs) are rapidly gaining recognition as critical mediators of inter-cellular communication during viral infections. To contribute to fill the gap in knowledge regarding the role of EVs in adenovirus infection, we used human adenovirus type 4 of species *Mastadenovirus exoticum* (HAdV-E4), a prevalent respiratory and ocular pathogen, and characterized the cargo and biological properties of EVs released by HAdV-E4-infected A549 lung epithelial cells at a pre-lytic stage of infection. Using immunocapture-based isolation and multi-omics approaches, we found that infection profoundly alters the EV uploaded proteome and small non-coding RNA repertoire. Mass spectrometry identified 268 proteins unique to EVs purified from infected cells (AdV-EVs), with enrichment in pathways supporting vesicle trafficking and viral protein translation, and importantly also a few virus-encoded proteins. Small RNA transcriptome analysis showed differential uploading in AdV-EVs of various small non-coding RNAs, including snoRNAs, as well as the presence of virus associated RNAs I and II. Notably, AdV-EVs contained viral genomic DNA and were capable of initiating productive infection upon delivery to naïve cells in the absence of detectable viral particles. Our data suggest that EVs released during HAdV-E4 infection may serve as vehicles for non-lytic viral dissemination and highlight their possible role in intra-host dissemination.

## INTRODUCTION

Extracellular vesicles (EVs) are a heterogeneous class of particles of various sizes delimited by a lipidic bilayer that are released by virtually all cell types and can be generated either in the cell internal compartments by the multivesicular body (exosomes), or by budding off the cell surface (ectosomes or microparticles) [1]. During their biogenesis, EVs acquire a variety of molecules that include membrane and cytosolic proteins, all classes of RNAs, and DNA [2, 3]. The specific EV composition varies based on the type of originating cell and can change depending on the cell response to external stimuli [4]. A fundamental property of EVs is the ability to transfer their cargo to neighboring or distant cells and thus affect their biological functions. EVs have gained recognition as important mediators of intercellular communication and play significant roles in a number of diseases, including infectious diseases [5]. In the context of viral infections, there is growing evidence that a number of viruses can utilize EVs as shuttles to transfer viral components, genetic material, or even complete virions, which in turns favors virus dissemination and increases the virus ability to evade the immune system [6]. Studies have shown that several non-enveloped viruses can take advantage of different EV biogenetic processes in order to acquire an envelope and be released non-lytically to the extracellular space. In particular, picornaviruses hepatitis A virus and enterovirus A71 exploit the endosomal sorting complexes required for transport (ESCRT)- mediated exosome biogenesis pathway [7, 8], while coxsackievirus B3 and rhesus rotavirus can be shed through microvesicles originating from the plasma membrane of infected cells [9, 10]. Interestingly, poliovirus, another picornavirus, can be encapsulated either in microvesicles or in autophagosome-derived vesicles for non-lytic exit [11].

Human adenoviruses (HAdVs) are non-enveloped, double-stranded DNA viruses in the genus *Mastadenovirus* of the *Adenoviridae* family and are grouped into seven species (HAdV-A through G). HAdVs are causative agents of respiratory diseases of variable severity (species B, C and E), conjunctivitis (species B, D and E), and gastroenteritis (species F and G) [12, 13]. While most infections are self-limited and associated with mild clinical presentations in immunocompetent hosts, they can be life-threatening in individuals with a compromised immune system, such as patients with primary immunodeficiencies and transplant recipients. In the latter, detected infections are most frequently the result of viral reactivation from latency and often become systemic affecting various organ systems [14]. At present, there are no approved antivirals specific for adenovirus and the only available vaccine targets types E4 and B7 and is licensed for use exclusively in military recruits entering basic training and other military personnel at high risk for HAdV infection [15].

So far, only a few studies have provided insight into the possible involvement of EVs in adenovirus infection. These studies, however, were carried out with a non-replicative HAdV-B3 [16] or with oncolytic HAdV-C5 based vectors [17–21] leaving opportunity to examine the characteristics and role of EVs in adenovirus infection using more pathophysiologically relevant experimental systems. In this study, we show that HAdV-E4 infection of lung epithelial cells elicits profound changes in the protein and RNA cargo composition of EVs isolated from infected cell supernatants at a pre-lytic time post infection. Furthermore, we found that EV-associated HAdV-E4 DNA can be transferred to naïve cells and initiate a productive infection.

## MATERIALS AND METHODS

### Cell culture

A549 human type II alveolar epithelial cells (ATCC CL-185) were grown in minimum essential medium (MEM; ThermoFisher Scientific, Waltham, MA, USA) supplemented with 8% heat-inactivated newborn calf serum (HI-NBCS; Rocky Mountain Biologicals, Missoula, MT, USA), 1.5 g/L sodium bicarbonate (MilliporeSigma, Burlington, MA, USA), 2mM L-glutamine, 100 U/mL penicillin, 100 µg/mL streptomycin) and 25 mM HEPES (all from ThermoFisher Scientific). For EV isolation, A549 cells were propagated in roller bottles in RPMI 1640 medium (ThermoFisher Scientific) containing 10% HI-NBCS, 1.05 g/L sodium bicarbonate, 25 mM HEPES, 100 U/mL penicillin and 100 µg/mL streptomycin, at 37 °C.

### Virus propagation and stock production

The prototype-like vaccine strain CL68578 of human adenovirus E4 (HAdV-E4) used in these studies was derived from the pVQ WT number 11183 genomic clone obtained from Advanced Bioscience Laboratories (ABL), Inc. as previously described [22] by transfection of A549 cells in culture. Stocks were produced at passage 4-5 and their infectious titers determined by plaque assay as described [22].

### Viral Infection for EV isolation

A549 cell monolayers in roller bottles were infected with HAdV-E4 at a multiplicity of infection of 10 PFUs/cell in Hanks Balanced Salt Solution (HBSS; ThermoFisher Scientific). After 2 hours absorption, cells were rinsed three times with PBS and replenished with MEM containing 1% EV-depleted HI-NBCS (by centrifugation at 120,000 x g for 16 hours at 4 °C in a Beckman Optima XL 100 ultracentrifuge equipped with a SW 32 Ti rotor), 1.05 g/L sodium bicarbonate, 25 mM HEPES, 100 units/mL penicillin and 100 µg/mL streptomycin. At 24 hours post-infection, conditioned medium from mock-infected or HAdV-E4-infected cells was harvested and centrifuged at 300 x g for 10’ at 4 °C, then at 10,000 x g for 30’ at 4 °C. The resulting clarified medium was concentrated by ultrafiltration with a 100 KDa cutoff regenerated cellulose membrane in a Centricon unit (MilliporeSigma), at 4 °C, until reduced to a volume of ∼ 2 mL. EVs were isolated from the concentrated medium with EV isolation kits Pan (human), or CD81 human (Cat # 130-111-572 and 130-110-914, Miltenyi Biotec, Auburn, CA, USA), according to the manufacturer’s instructions. Purified EVs were either processed immediately, or stored at −80 °C for further analysis.

### Western blot

EV proteins were obtained by lysis with RIPA buffer containing protease inhibitors (MilliporeSigma, Burlington, MA) and quantified with Micro BCA protein assay kit (ThermoFisher Scientific). Protein extracts were separated on a 4-12% Bolt Bis-Tris Plus Mini Protein Gel (ThermoFisher Scientific), subsequently blotted onto a 0.2 µm PVDF membrane (ThermoFisher Scientific) and then blocked with 5% BSA (MilliporeSigma) at 37 °C for 1 hour. Incubation with primary antibodies anti-huCD9 (mouse mAb, clone MM2/57, cat #CBL 162, MilliporeSigma), anti-huCD63 (Cat# 556019, BD Pharmingen, Franklin Lakes, NJ, USA), anti-CD81 (mouse mAb cat # SC-166029, Santa Cruz Biotechnology, Dallas, TX, USA), anti-Alix (rabbit mAb E6P9B cat #92880 Cell Signaling Technology, Danvers, MA, USA) and anti GRP94 (rabbit mAb D6X2Q cat# 20292, Cell Signaling Technology, Danvers, MA, USA) was carried out at 4 °C overnight. Following incubation with appropriate HRP-conjugated secondary antibodies for 1 hour at room temperature, immunoreactive proteins were visualized using SuperSignal West Femto Maximum Sensitivity Substrate (ThermoFisher Scientific). Images were acquired with a ChemiDoc MP Imaging System (Bio-Rad, Hercules, CA, USA), and analyzed with Image Lab software 6.1.0 (Bio-Rad).

### Transmission electron microscopy (TEM)

TEM was performed at the Electron Microscopy core facility at the University of New Mexico (UNM). Aliquots of 10 μl of EV preparations were fixed with 1% glutaraldehyde overnight at 4°C and transferred to UNM for negative staining with uranyl acetate, and imaging in a Hitachi HT7700 microscope equipped with an AMT XR16M 16-megapixel digital camera. Ten to fifteen images were acquired from each grid at various magnifications.

### Nanoparticle tracking analysis (NTA)

NTA was performed by Alpha Nano Tech, LLC (Research Triangle Park, NC, USA). Samples were fixed with 1% paraformaldehyde before analysis to inactivate any potentially infectious particles. Samples were incubated with 50 µg/mL of the lipophilic membrane dye CMDR (ThermoFisher Scientific) for 30 minutes in the dark at room temperature, diluted with high purity water, and analyzed in a Zetaview Quatt instrument (Particle Metrix, Ammersee, Germany) in both scatter and fluorescent mode.

### Flow cytometry analysis of EV surface markers

The characterization of surface markers was carried out on EVs isolated with CD81 EV isolation kit (Miltenyi Biotec) using the MACSPlex Exosome kit (Miltenyi Biotec) according to the manufacturer’s instructions. Cytofluorimetric analyses were performed on a BD FACSCelesta flow cytometer equipped with BD FACSDiva software V 9.0 (BD Biosciences, Franklin Lakes, NJ, USA). Data were normalized on the average of CD9 and CD63 median fluorescence intensities. The protein-protein interaction network analysis was performed with STRING (https://string-db.org).

### Proteome profiling

EV proteins were obtained by lysis with RIPA buffer containing protease inhibitors (MilliporeSigma). For the immunoprecipitation experiments, 5 µg of EV lysates were incubated with 10 µL of heat inactivated anti-HAdV-E4 rabbit antiserum (S-1001, a generous gift from the Viral and Rickettsial Disease Laboratory, California Department of Health Services) overnight at 4°C. Immunocomplexes were recovered using MagnaBind protein G magnetic beads (ThermoFisher Scientific). At IDeA National Resource for Quantitative Proteomics total protein from each sample was reduced, alkylated, and digested using single-pot, solid-phase-enhanced sample preparation [23] with sequencing grade modified porcine trypsin (Promega, Fitchburg, WI, USA). Tryptic peptides were then separated by reverse phase XSelect CSH C18 2.5 um resin (Waters, Milford, MA, USA) on an in-line 150 x 0.075 mm column using an UltiMate 3000 RSLCnano system (ThermoFisher Scientific). Peptides were eluted with a mixture of 0.1% formic acid, 0.5% acetonitrile (buffer A) and 0.1% formic acid, 99.9% acetonitrile (buffer B), using a 60 min gradient from 98:2 to 65:35 buffer A:B ratio. Eluted peptides were ionized by electrospray (2.2 kV) followed by mass spectrometric analysis on an Orbitrap Exploris 480 mass spectrometer (ThermoFisher Scientific). To assemble a chromatogram library, six gas-phase fractions were acquired on the Orbitrap Exploris with 4 m/z DIA spectra (4 m/z precursor isolation windows at 30,000 resolution, normalized AGC target 100%, maximum inject time 66 ms) using a staggered window pattern from narrow mass ranges using optimized window placements. Precursor spectra were acquired after each DIA duty cycle, spanning the m/z range of the gas-phase fraction (i.e. 496-602 m/z, 60,000 resolution, normalized AGC target 100%, maximum injection time 50 ms). For wide-window acquisitions, the Orbitrap Exploris was configured to acquire a precursor scan (385-1015 m/z, 60,000 resolution, normalized AGC target 100%, maximum injection time 50 ms) followed by 50x 12 m/z DIA spectra (12 m/z precursor isolation windows at 15,000 resolution, normalized AGC target 100%, maximum injection time 33 ms) using a staggered window pattern with optimized window placements. Precursor spectra were acquired after each DIA duty cycle. Following data acquisition, data were searched using Spectronaut (Biognosys AG, Zurich, Switzerland) v 18.5 against the UniProt Homo sapiens plus human adenovirus E serotype 4 database (April 2023) using the directDIA method with an identification precursor and protein q-value cutoff of 1%, generate decoys set to true, the protein inference workflow set to maxLFQ, inference algorithm set to IDPicker, quantity level set to MS2, cross-run normalization set to false, and the protein grouping quantification set to median peptide and precursor quantity. Protein MS2 intensity values were assessed for quality using ProteiNorm [24]. The data was normalized using VSN [25] and analyzed using proteoDA to perform statistical analysis using Linear Models for Microarray Data (limma) with empirical Bayes (eBayes) smoothing to the standard errors [26, 27]. Proteins with an FDR adjusted p-value < 0.05 and a fold change > 2 were considered significant. Gene ontology (GO) and network analysis were performed using ClueGO v. 2.5.8 and Cytoscape v. 3.9.1. EV protein databases ExoCarta (http://www.exocarta.org) [28] and Vesiclepedia (http://microvesicles.org) [29] were used as a reference for analysis of identified proteins.

### Characterization of small RNA cargo

Total RNA was isolated from EVs with TRIzol LS reagent (ThermoFisher Scientific) and miRNeasy Micro Kit (Qiagen, Germantown, MD, USA). Residual DNA was removed by in-column treatment with RNase-Free DNase Set (Qiagen). Sample quality was assessed by High Sensitivity RNA Tapestation (Agilent Technologies Inc., Santa Clara, CA, USA) and yield quantified by Qubit RNA HS assay (ThermoFisher Scientific). Sequencing services were contracted from CD Genomics (Shirley, NY, USA). Library preparation was performed with SMARTer Small RNA (Takara Bio USA Inc., San Jose, CA, USA) following manufacturer’s instructions. Samples were pooled and sequenced on an Illumina Novaseq S4 sequencer for 150 bp read length in paired-end mode. Reads were mapped to the reference genomes by HISAT2, and htseq-count software was used for quantification. Annotation sources for the small RNAs were miRBase v. 22 for miRNAs, piRNABank, piRBase and piRNACluster for piRNAs, and GENCODE release 27 for snRNAs and snoRNAs. The RNAInter database, a curated repository of experimental RNA interactions [30], was used to identify potential interactors for snoRNAs. Only interactions with confidence score ≥ 0.25 were considered. The human adenovirus E, complete genome (GenBank accession #NC_003266.2) was used for virus associated (VA) RNA identification. DESeq2 software was used for differential abundance analysis. GO and network analysis were performed using Cytoscape v. 3.9.1 and ClueGO v. 2.5.8. For validation of VA RNA detection, 10 ng of total RNA isolated as described above were reverse transcribed with SuperScript III First-Strand Synthesis System and random primers (ThermoFisher Scientific) according to the manufacturer’s instructions. cDNA was amplified using PowerUP SYBR Green Master Mix (ThermoFisher Scientific) with the following primers and concentrations: VA RNA_I_ FW 5’-CTAAGCGAACGGGTTGGGCTG-3’, 400 nM, rv 5’-CCAGTACCACGTTAGCTGCGG-3’, 200 nM; VA RNA_II_ FW 5’-AGAATCGCCAGGGTTGCGTTG-3’, 400nM, RV 5’-TTGGAAACGACGGGGCAGC-3’, 400 nM; ACTB (β-actin) FW 5’-TCCTCCTGAGCGCAAGTACTC-3’, 200 nM, RV 5’-CGGACTCGTCATACTCCTGCT-3’, 200 nM. Amplification was carried out in a QuantStudio 5 Real-Time PCR System (Thermo Fisher Scientific) and the following cycling conditions: 95.0°C for 10’ followed by 40 cycles of 95.0°C for 15’’ and 60.0°C for 1’. Data analysis was performed with the QuantStudio Design & Analysis Software v1.5.2 (Thermo Fisher Scientific).

### Infectivity and neutralization assays

Infectivity of EV preparations was assessed by plaque assay in 12 well-plates. Infectious EV titers were expressed as plaque forming units (PFUs)/mL. Neutralization assays were carried out using heat-inactivated reference rabbit antiserum S-1001 to HAdV-E4 (prototype strain RI-67), a generous gift from Dr, Shigeo Yagi, California Department of Public Health. Two-fold serial dilutions of serum were incubated with approximately 20 - 30 PFUs of purified HAdV-E4 or 20 - 30 PFUs of purified EVs for 1 hour at 37 °C and then neutralization mixes were delivered onto A549 cell monolayers in triplicate. After adsorption for 1 hour at 37 °C, monolayers were overlaid with 0.55% low melt agarose in MEM.

### Triton, Proteinase K and DNase treatments of EV preparations

EVs were lysed in the presence of 1% Triton X-100 (Bio-Rad) on ice for 10’, where indicated. EVs were then incubated with 10 mg/mL proteinase K (Bioline, Taunton, MA, USA) for 2 h at 37 °C. Proteinase K activity was stopped by addition of 1 mM PMSF (MilliporeSigma), followed (or not) by digestion with 10 U of DNase I (ThermoFisher Scientific) for 30’ at 37 °C. Treated samples were processed for qPCR and plaque assay as described below.

### Quantitative PCR (qPCR) detection of viral DNA

DNA was extracted from EV preparations derived from infected cells with DNeasy Blood and Tissue Kit (Qiagen), according to manufacturer’s instructions. 9 µL of isolated DNA were used per each reaction. DNA amplification was carried out with TaqMan Fast Advanced Master Mix (Thermo Fisher Scientific) in the presence of 250 nM probe and 900 nM of each primer in 20 µL of reaction mix, using a QuantStudio 5 Real-Time PCR System (ThermoFisher Scientific). Cycling conditions were as follows: 95.0°C for 20’’ followed by 40 cycles of 95.0°C for 1’’ and 60.0°C for 20’’. Probe and primers were modified from [31], and obtained from Integrated DNA Technologies (Coralville, IA, USA): FW 5’-GCCCCAGTGGTCTTACATGCACATC-3’; RV 5’-GCCACCGTGGGGTTTCTAAACTT-3’; probe 5’-/56-FAM/TGCACCAGACCCGGGCTCAGGTACTCCGA/36-TAMSp/-3’. DNA isolated from purified HAdV-E4 viral particles was quantified with Qubit DNA HS assay (ThermoFisher Scientific) and used to generate a standard curve ranging from 1X10^0^ to 1×10^9^ genome copies per reaction. The lower detection limit of the developed assay was 200 genome copies per reaction. All the reactions were carried out in duplicate. Data analysis was performed with the QuantStudio Design & Analysis Software v1.5.2 (ThermoFisher Scientific).

## RESULTS

### Characterization of EVs from mock- and HAdV-E4-infected A549 cell supernatants

The population of EVs released from A549 cells in the early non-lytic stages of HAdV-E4 infection was characterized using several approaches. EVs were isolated from the supernatant of mock-infected (Ctr-EVs) or HAdV-E4-infected (AdV-EVs) A549 cell monolayers at 24 hours post-infection (hpi), when a clear and wide-spread cytopathic effect was visible, but before cell lysis and consequent release of viral progeny. As shown in Figure 1A, cells were still attached and viable. To minimize the likelihood of presence of contaminating viral particles in the EV preparations, we opted for an immunocapture-based EV isolation method in which EVs are isolated from the cell culture supernatants by using magnetic beads conjugated to antibodies against the EV markers tetraspanins CD9, CD63, and CD81[32]. The purity, size distribution, and concentration of the isolated EVs was assessed by western blot, transmission electron microscopy (TEM), and nanoparticle tracking analysis (NTA) to obtain basic information on the preparations following the latest recommendations of the International Society for Extracellular Vesicles (ISEV) [1]. As shown in Figure 1B, both Ctr-EVs and AdV-EVs preps exhibited a marked enrichment of EV markers CD63, CD9, CD81 and Alix compared to their respective cells of origin. Absence of the endoplasmic reticulum marker GRP94 in EV lysates indicated negligible source cellular compartment contamination. The analysis of TEM images from Ctr-EVs and AdV-EVs revealed the presence of rounded particles with a diameter ranging from 50 to 150 nm and, importantly, absence of contaminating viral particles in EV preps purified from HAdV-E4-infected cell supernatants. (Fig. 1C). The ability to visualize and discriminate HAdV particles from EVs was verified by examining a sample of purified HAdV-E4 particles, which displayed the characteristic icosahedral shape and a diameter of about 70 nm (Fig. 1D). To better characterize EV size distribution and measure particle concentrations, both Ctr-EVs and AdV-EVs were subjected to NTA. To specifically detect signals from the EVs and exclude from the analysis the magnetic beads to which they were attached as a result of the immunomagnetic purification procedure, EV membranes were labelled with the lipophilic fluorescent dye CMDR and signals were acquired in both scatter and fluorescent mode. A control performed with magnetic beads alone demonstrated absence of fluorescent signal after CMDR labelling (Fig. 1E). Ctr-EVs and AdV-EVs showed a similar size distribution profile, with a peak at around 170 nm, and the number of particles obtained in the two preparations was also similar (Table 1 and Fig. 1E). The detected particle size was slightly higher than expected, but this can be explained by the fact that each magnetic bead can capture multiple EVs, as readily visible in the TEM images.

**Figure 1.**
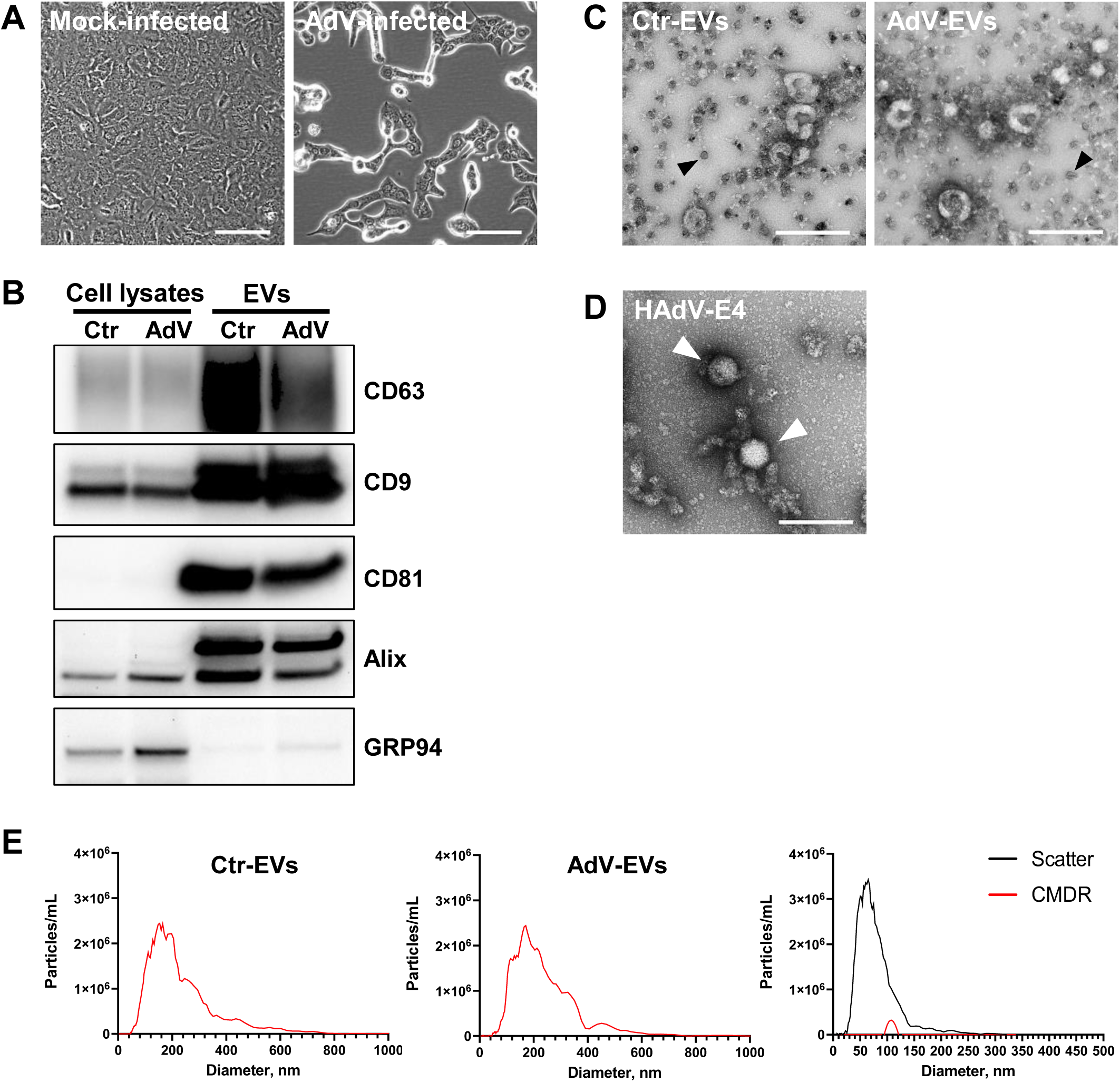
Characterization of the EV populations isolated from mock- and HAdV-E4-infected A549 cells. **A.** Phase contrast images of A549 cells showing the magnitude of cytopathic effect at 24 hours post-infection. Cells were infected at MOI = 10. Scale bars = 100 µm. **B.** Representative Western blot analysis showing enrichment of the EV markers CD63, CD9, CD81, Alix in EVs-Ctr and EVs-AdV, compared to their cells of origin (cell lysates). The ER marker GRP94 was used to assess the presence of non-EV contaminants. Images are representative of three different experiments. **C.** TEM images of EVs isolated by magnetic immunocapture form the supernatants of mock-infected cells (Ctr-EVs) or HAdV-E4-infected (AdV-EVs) A549 cells. The magnetic beads are recognizable by their opaque appearance and uniform size of 50 nm (black arrows). Scale bars = 200 nm**. D.** TEM image of purified HAdV-E4 particles (white arrows). Scale bar = 200 nm. **E.** Fluorescent nanoparticle tracking analysis of Ctr-EVs (left panel), AdV-EVs (middle panel), and magnetic beads alone (right panel).

**Table 1:** Vesicle size distribution and concentration of purified Ctr- EVs and AdV- EVs.

| | Median Size ± SD | Concentration (particles/mL ± SD) |
| --- | --- | --- |
| Ctr-EVs | 176 ± 18 nm | $1.45 \times 10^{11} \pm 5.00 \times 10^9$ |
| AdV-EVs | 165 ± 5 nm | $1.17 \times 10^{11} \pm 9.75 \times 10^9$ |

### Characterization of AdV-EVs and Ctr-EVs surface markers

To better characterize the protein composition of the EV membranes, Ctr-EVs and AdV-EVs were screened for the presence of 37 different proteins by flow cytometry. Consistent with their epithelial origin, both EV populations expressed the epithelial marker EpCAM (CD326). The other proteins detected were CD40, CD44, ROR1, HLA-ABC, CD24 and ITGB1 (Integrin beta-1, also known as CD29), which were less abundant in AdV-EVs compared to Ctr-EVs (Fig. 2A). A functional enrichment analysis conducted by interrogating the STRING protein interaction database [33] indicated that EpCAM, CD40, CD44, CD24 and ITGB1 are part of a functional protein association network (enrichment p-value = 1.37 x 10^-5^), which includes a direct interaction between CD44 and ITGB1 (Fig. 2B). Interestingly, CD40, a member of the TNF-receptor superfamily that mediates humoral and adaptive immune responses in the lung [34] was the surface protein with the lowest abundance in AdV-EVs compared to Ctr-EVs.

**Figure 2.**
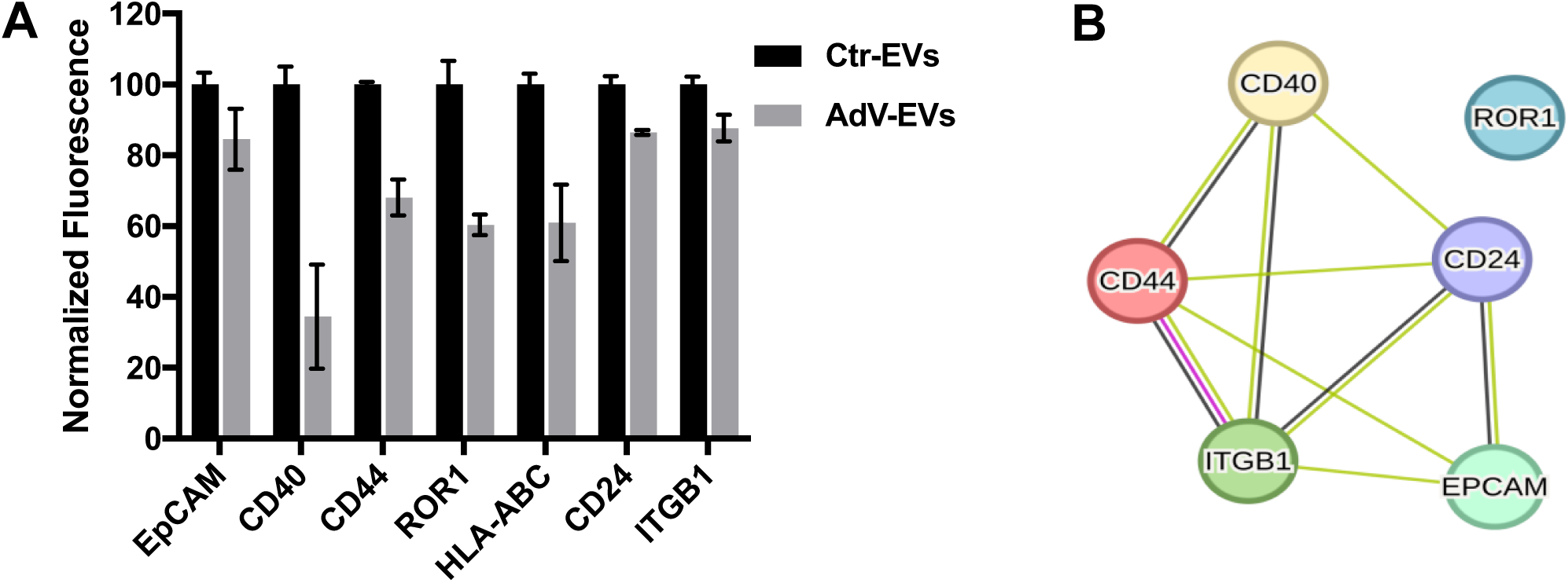
Characterization of surface markers of AdV-EVs and Ctr-EVs. A. MACSPlex flow cytometry detection of surface markers. Data represent ratios of normalized median fluorescent intensity (MFI) in AdV-EVs over Ctr-EVs. The mean values from of two independent experiments ± SEM are shown. B. STRING protein-protein interaction network analysis of the proteins shown in (A). Yellow lines represent text mining evidence, black lines represent co-expression evidence, purple line represents experimental evidence.

### Proteomics profiling of AdV-EVs and Ctr-EVs

To determine whether HAdV-E4 infection induces alterations in the protein composition of EVs secreted by A549 cells, matched biological replicates of AdV-EVs (n=5) and Ctr-EVs (n=5) were isolated as described above and analyzed by data-independent acquisition mass spectrometry (DIA-MS). Proteins with an average spectral count > 2 for each group were considered. A total of 1837 proteins were identified in Ctr-EVs and 2083 in AdV-EVs. Of these, 1815 proteins (86.2%) were in common between Ctr-EVs and AdV-EVs; 22 proteins (1.0%) were exclusively present in Ctr-EVs, while 268 (12.7%) were exclusively present in AdV-EVs (Fig. 3A). Proteins commonly found in EVs were detected in both the Ctr-EVs and AdV-EVs datasets. Among the proteins identified in both Ctr-EVs and AdV-EVs, 23 were present only in the ExoCarta list, 23 were present only in the Vesiclepedia list, and 67 proteins were present in both the ExoCarta and Vesiclepedia lists (Fig. 3B). To obtain more specific information regarding the protein composition of EVs derived from HAdV-E4-infected A549 cells, we performed a gene ontology (GO) and network analysis on the 268 proteins that were exclusive to AdV-EVs. Significantly enriched GO terms were grouped in an annotation network based on the presence of similar or common proteins. The resulting network was composed of 4 major components, in which the most representative terms were endosomal transport, vesicle targeting, vesicle organization, vesicle fusion, and cytoplasmic translation (Fig. 3C). Since these proteins were not detected in Ctr-EVs, their presence in AdV-EVs may be indicative of increased vesicle trafficking and increased viral protein translation in the originating cells as a consequence of the ongoing viral infection. To compare the relative protein abundance between AdV-EVs and Ctr-EVs, normalized exclusive intensities were used for differential analysis. A total of 326 proteins were differentially abundant (|log2 fold change| > 1, pval < 0.05). Of these, 107 were more abundant in AdV-EVs and 219 were less abundant in AdV-EVs compared to Ctr-EVs, as shown in the volcano plot in Fig. 4A. A hierarchical clustering of the differentially abundant proteins was generated and the corresponding heatmap shows that Ctr-EVs and AdV-EVs cluster separately (Fig. 4B). The top 10 proteins with increased or decreased abundance in AdV-EVs compared to Ctr-EVs are shown in Table 2. Differentially abundant proteins were further investigated by GO analysis. The most enriched biological processes for the proteins with increased abundance in AdV-EVs included GO terms representative of protein translation in response to viral infections, cytoskeletal remodeling, interactions with the extracellular matrix, and mTOR and ErbB2 signaling pathways (Fig. 4C). Interestingly, adenovirus proteins E4-ORF1 and E4-ORF4 can activate mTOR signaling to promote protein translation and thus facilitate viral replication [35], and E1A can affect ErbB2 expression and inhibit cell proliferation [36]. Proteins with decreased abundance in AdV-EVs showed enrichment for terms related to the Toll-like receptor signaling, tumor necrosis factor (TNF), interleukin-mediated inflammatory response, and activation of the complement pathway, which are all components of the innate antiviral immune response [37, 38] (Fig. 4D). The differential abundance analysis confirmed that CD40, which showed reduced surface expression in AdV-EVs (Fig. 2), was less abundant in AdV-EVs in terms of total proteins, and contributed to the TNFR/NF-kB-related pathways illustrated in Fig. 4D. The proteomics dataset was also mined for the presence of HAdV-E4-encoded proteins. A set of 8 adenoviral proteins were detected in AdV-EVs. Of these, 6 are structural components of the virion and two, E2A-DBP (DNA Binding Protein) and L4-100K, are non-structural proteins (Fig. 5). DBP plays a critical role in the AdV life cycle controlling viral DNA replication and transcription, and mRNA stability [39]. L4-100K, also known as shutoff protein, blocks the translation of host proteins, while promoting the translation of viral proteins [40].

**Figure 3.**
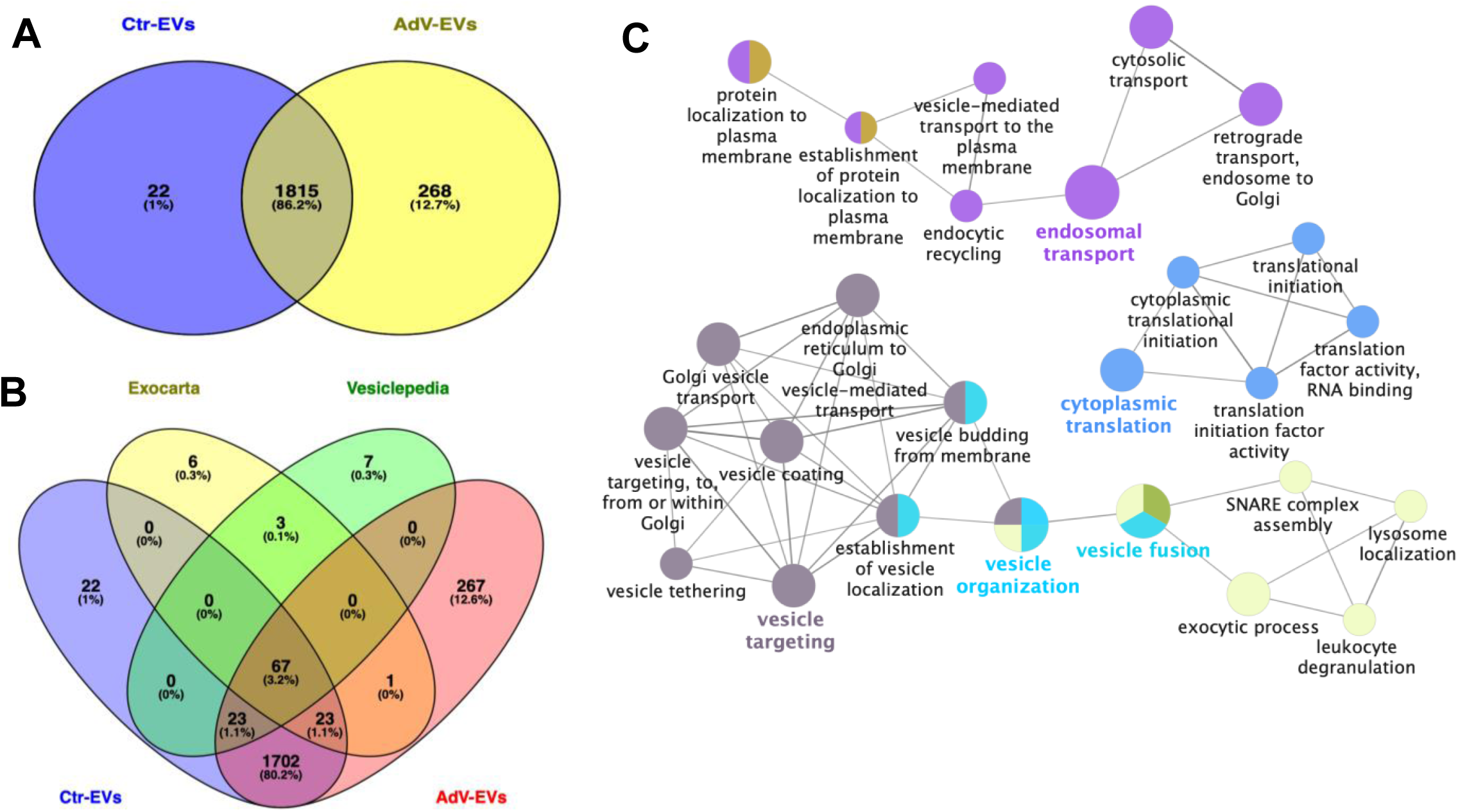
Proteome profiling of Ctr-EVs and AdV-EVs. A. 2-way Venn diagram of proteins identified in Ctr-EVs and AdV-EVs with spectral counts ≥ 2 in 5 biological replicates. 22 and 268 proteins were unique to Ctr-EVs and AdV-EVs, respectively. B. 4-way Venn diagram showing the intersection between proteins in (A) and the top-100 EV proteins included in the Exocarta and Vesiclepedia repositories. C. GO and network analysis of proteins found exclusively in AdV-EVs. Nodes represent GO terms with adjusted p value < 0.05 after Benjamini-Hochberg correction. Ontology sources were Biological Process, Reactome Pathways and Immune System.

**Figure 4:**
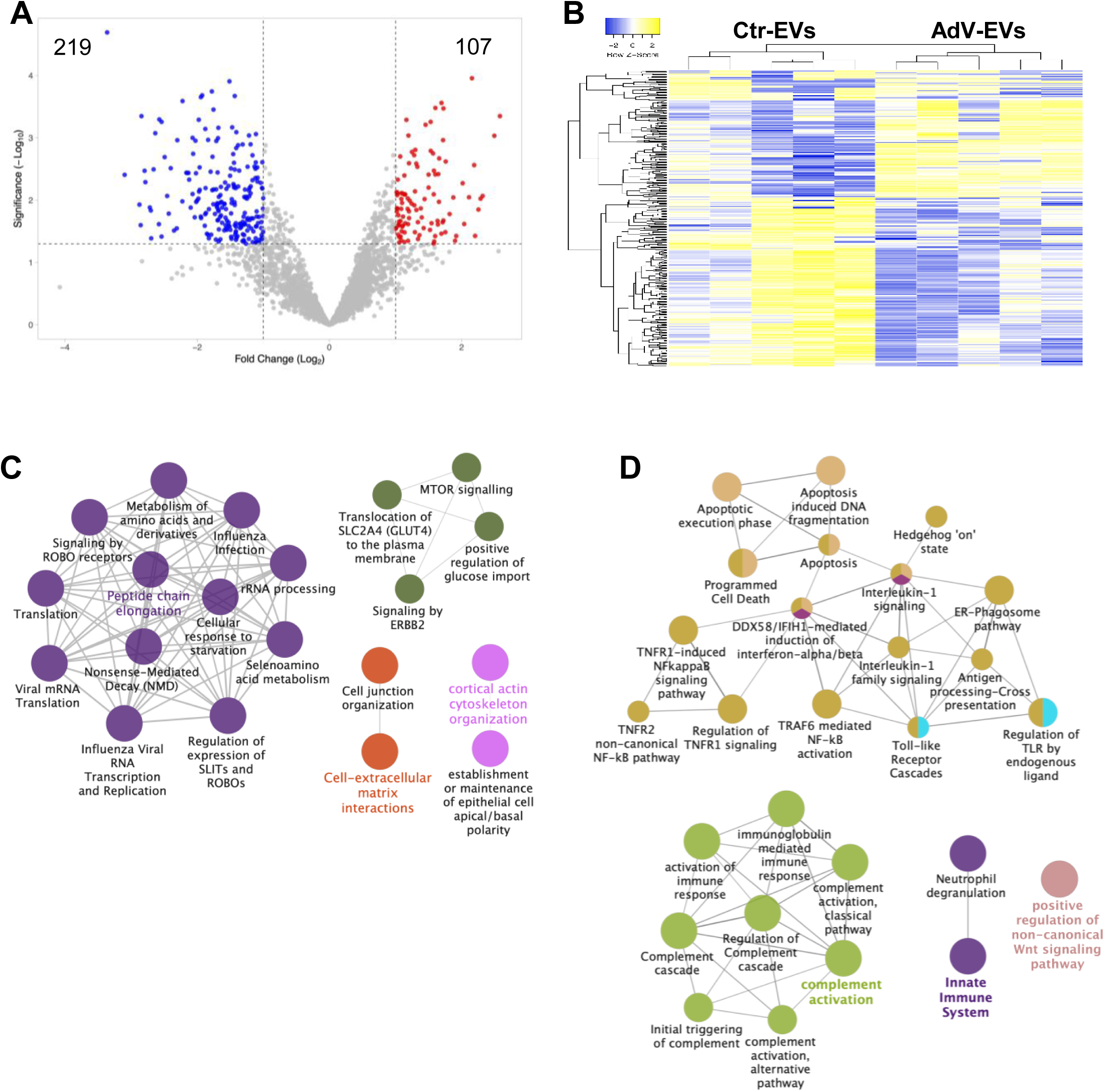
Functional analysis of differentially abundant proteins in AdV-EVs versus Ctr-EVs. **A.** Volcano plot of differentially abundant proteins in AdV-EVs versus Ctr-EVs. 107 proteins were more abundant in AdV-EVs (log2FC > 1, Pval < 0.05), and 219 proteins were less abundant in AdV-EVs (log2FC < 1, Pval < 0.05). **B.** Hierarchical clustering heatmap of differentially abundant proteins. Data from 5 biological replicates are shown. Yellow indicates increased abundance; blue indicates decreased abundance. **C,D**. GO and network analysis of more abundant (C) and less abundant (D) proteins in AdV-EVs. Nodes represent GO terms with adjusted p value < 0.05 after Benjamini-Hochberg correction. Ontology sources were Biological Process, Reactome Pathways and Immune System.

**Figure 5:**
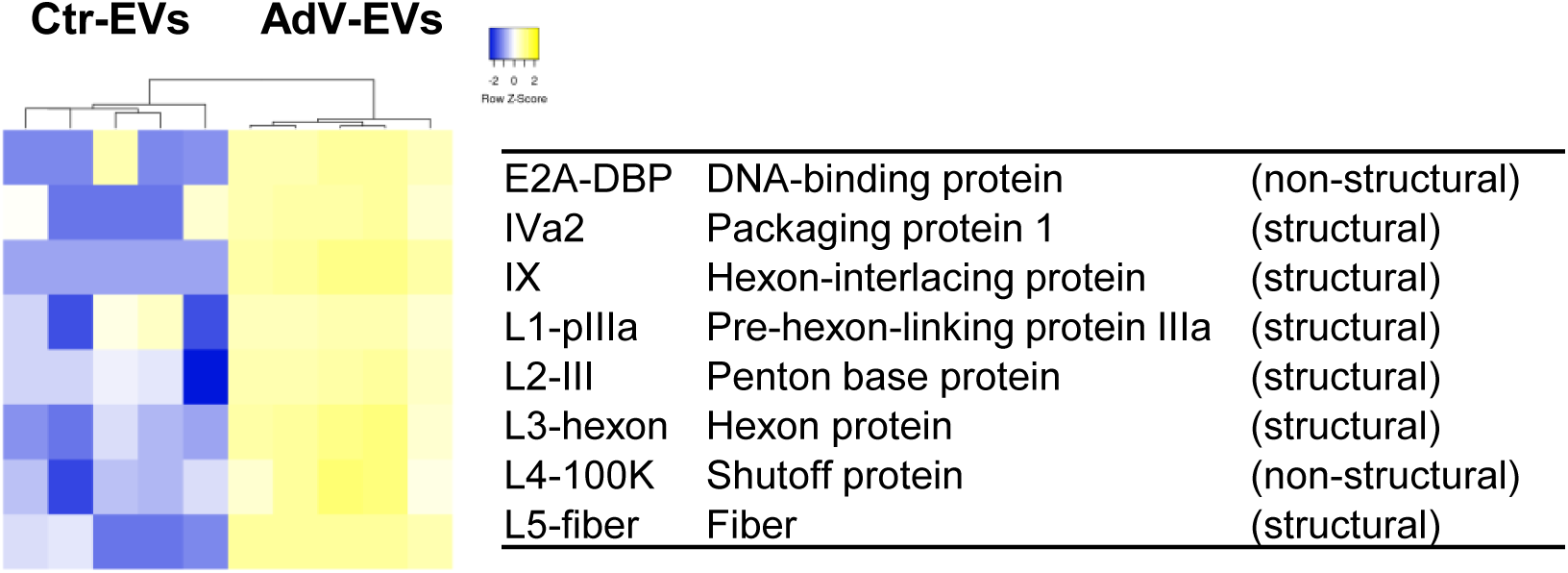
Detection of adenovirus proteins in EVs-AdV. Hierarchical clustering heatmap of 6 structural and two non-structural HAdV-E4 proteins identified in AdV-EVs. Data from 5 biological replicates are shown. Yellow indicates increased abundance; blue indicates decreased abundance.

**Table 2:** Top 10 proteins with increased or decreased abundance in AdV-EVs compared to Ctr-EVs.

| Increased abundance |  | Decreased abundance |  |
| --- | --- | --- | --- |
| Protein name | log2 fold change | Protein name | log2 fold change |
| FUS | 2.58 | SLITRK6 | -3.36 |
| SLC20A1 | 2.49 | UBB;UBC | -3.10 |
| RPS6 | 2.32 | TRAF2 | -2.87 |
| RPL7 | 2.29 | KRT19 | -2.86 |
| RPL15 | 2.25 | PDGFA | -2.84 |
| COMT | 2.20 | DKK1 | -2.80 |
| RPLP0 | 2.19 | CALML5 | -2.78 |
| DTNB | 2.16 | KRT6B | -2.72 |
| RPL4 | 2.11 | RSPO3 | -2.70 |
| KLHL13 | 2.04 | IFT81 | -2.70 |

### Small RNA content of AdV-EVs and Ctr-EVs

EVs can carry a spectrum of RNA molecules that encompass protein-coding RNAs and non-coding RNAs. Among the non-coding RNAs found in EVs, the small RNA class includes microRNAs (miRNAs), small nucleolar RNA (snoRNAs), small nuclear RNAs (snRNAs), piwi-interacting RNAs (piRNAs), transfer RNA (tRNAs), and ribosomal RNAs (rRNAs) [2]. To characterize the small RNA cargo of EVs released during the early non-lytic stages of HAdV-E4 infection, we performed small RNA sequencing of matched Ctr-EVs (n = 3) and AdV-EVs (n = 3) collected at 24 hpi. Data analysis identified a total of 813 and 806 unique transcripts with average counts >=2 in Ctr-EVs and AdV-EVs, respectively. The most represented class of small RNAs in both Ctr-EVs and AdV-EVs was piRNAs, followed by snoRNAs, snRNAs and miRNAs (Table 3). The relative abundance of each small RNA class in terms of mapped reads was also analyzed (Fig. 6A). snRNAs were the most abundant class in both Ctr-EVs and AdV-EVs, accounting for 53.5% and 49.1% of total mapped reads, respectively. piRNAs, which were the second most abundant class in Ctr-EVs, showed a drastic reduction in AdV-EVs, while miRNAs and especially snoRNAs showed increased abundance in AdV-EVs. The differential abundance analysis performed comparing AdV-EVs versus Ctr-EVs (|log2 fold change| > 1, adjusted pval < 0.05), confirmed the general increased transcript presence in the cargo of AdV-EVs, especially for snoRNAs, and identified 70 snoRNAs, 4 piRNAs and 3 miRNA precursors as upregulated, while 3 snRNAs were downregulated (Table 4 and Fig. 6B). Table 5 shows the 10 most upregulated and the 3 down-regulated small non-coding transcripts in AdV-EVs. The hierarchical clustering heatmap generated with the differentially uploaded transcripts shows that Ctr-EVs and AdV-EVs cluster separately (Fig. 6C). Because of the growing evidence of a role of certain classes of snoRNAs in viral infections [41], the RNAInter database was interrogated to identify potential interactors for the 70 upregulated snoRNAs, focusing in particular on RNA binding proteins (RBPs) as possible effectors. Among the total 376 predicted interactors, 40 were classified as RBPs. The two top interactors were METTL3 and FMR1, showing potential binding to 30 and 29 different snoRNAs, respectively. Interestingly, METTL3 is a cellular m^6^A writer enzyme recently shown to control the efficiency of HAdV-5 late gene transcript splicing [42], and FXR1 is a cellular RBP recently described as a novel m^6^A reader that regulates AdV late mRNA transcript stability [43].

**Figure 6:**
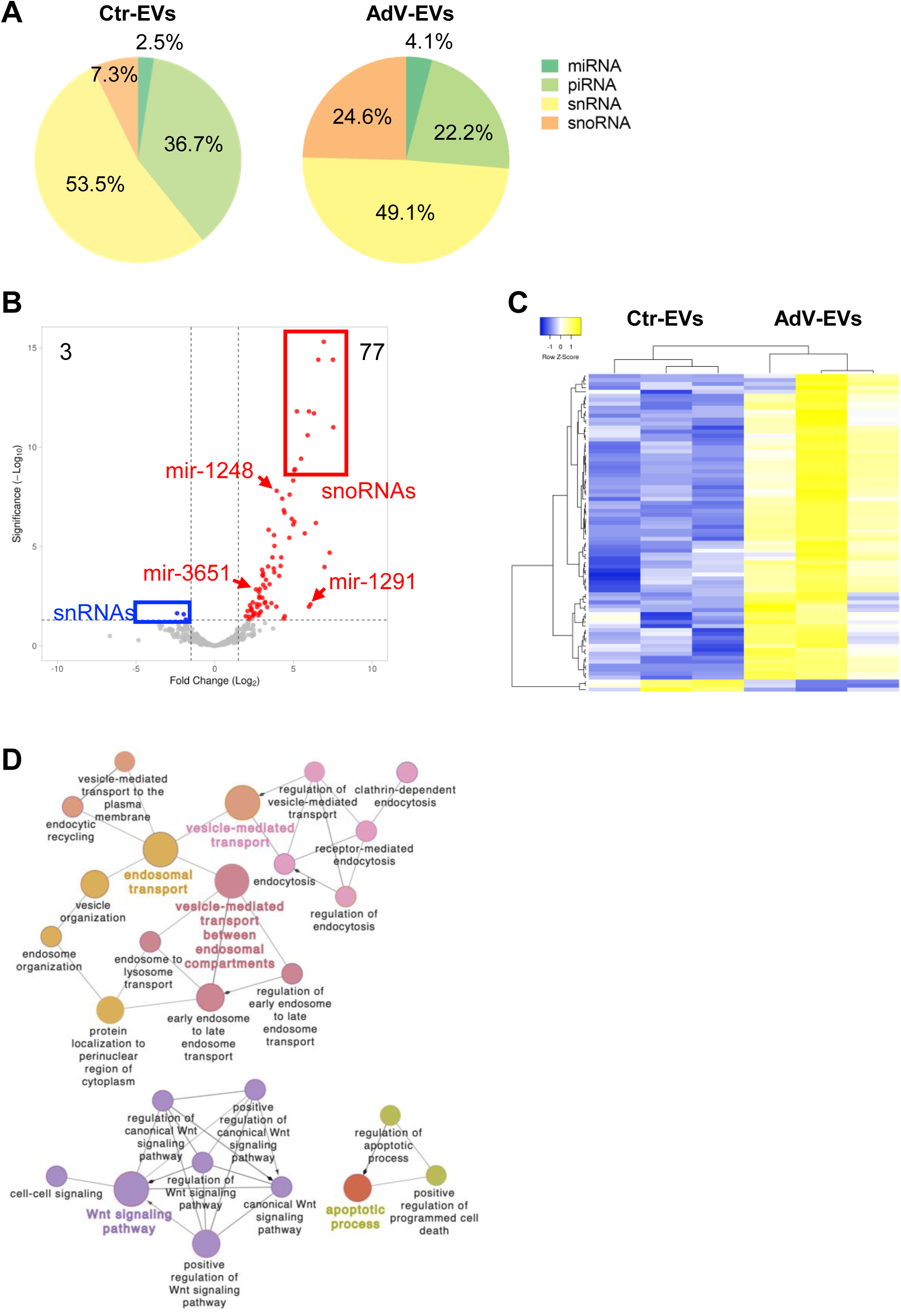
Analysis of the small RNA transcriptome of Ctr-EVs and AdV-EVs. **A.** Relative transcript abundance for each class of small RNAs in Ctr-EVs and AdV-EVs. **B.** Volcano plot of differentially uploaded small RNAs in AdV-EVs versus Ctr-EVs. 77 small RNAs were up-regulated in AdV-EVs (log2FC > 1, adjusted Pval < 0.05), and 3 small RNAs were down-regulated in AdV-EVs (log2FC < 1, adjusted Pval < 0.05). **C.** Hierarchical clustering heatmap of differentially uploaded small RNAs. Data from 3 biological replicates are shown. Yellow indicates increased presence; blue indicates decreased presence. **D.** GO and network analysis of mature miRNAs exclusively found in AdV-EVs. Nodes represent GO terms with adjusted p value < 0.05 after Benjamini-Hochberg correction. Ontology sources were Biological Process, Reactome Pathways and Immune System.

**Table 3:** Number of unique small non-coding RNA transcripts in Ctr- and AdV-EVs grouped by class.

| Class | Ctr- EVs | AdV- EVs |
| --- | --- | --- |
| miRNA (precursors) | 10 | 13 |
| miRNA (mature) | 40 | 28 |
| piRNA | 375 | 378 |
| snRNA | 124 | 107 |
| snoRNA | 264 | 280 |
| Total unique transcripts | 813 | 806 |

**Table 4:** Number of small non-coding RNA transcripts with differential relative abundance in AdV-EVs grouped by class.

| **Class** | **Higher abundance** | **Lower abundance** |
| --- | --- | --- |
| miRNA(p) | 3 | 0 |
| piRNA | 4 | 0 |
| snRNA | 0 | 3 |
| snoRNA | 70 | 0 |
| Total | 77 | 3 |

**Table 5:** Top transcripts with higher abundance and lower abundance in AdV-EVs compared Ctr-EVs.

| **Up-regulated** |  | **Down-regulated** |  |
| --- | --- | --- | --- |
| **Gene ID** | **log<sub>2</sub> fold change** | **Gene ID** | **log<sub>2</sub> fold change** |
| SNORA41 | 7.53 | RNU6ATAC2P | -2.39 |
| SNORA74A | 7.51 | RNU6ATAC | -1.97 |
| SNORA22 | 7.30 | RNVU1-1 | -1.84 |
| SNORA22C | 6.97 |  |  |
| SNORA50 | 6.92 |  |  |
| SNORA50C | 6.58 |  |  |
| SNORA50A | 6.43 |  |  |
| SNORA78 | 6.30 |  |  |
| hsa-mir-1291 | 6.08 |  |  |
| SNORA22B | 5.99 |  |  |

Among the 40 mature miRNAs identified in the dataset, 4 were exclusively detected in AdV-EVs, namely let-7f-5p, miR-93-5p, miR-7704 and miR-7706. To obtain predictive insights about the possible effect of EV-mediated delivery of these miRNAs to recipient naïve cells, we interrogated three different databases of miRNA-target interactions: TargetScan, microT-CDS and miRDB. The predicted target genes that were present in all three databases (intersection) were used for subsequent analyses. The resulting gene list was further filtered by using a publicly available expression dataset from The Genotype-Tissue Expression (GTEx) Project (https://www.gtexportal.org) consisting of 515 human lung tissue samples. From this dataset, only genes that had a median expression above a threshold of 3 transcript per million (TPM) were considered. The final gene list comprised 979 unique target genes, of which 388 were predicted targets of let-7f-5p, 626 of miR-93-5p, 5 of miR-7004, 15 of miR-7006. To identify biological pathways that could be potentially inhibited by the 4 miRNAs when delivered to recipient cells, the predicted target genes were analyzed by GO and the results visualized in an interaction network (Fig. 6D). The analysis showed enrichment for terms related to endosomal transport, vesicle trafficking, Wnt signaling pathway and apoptotic process. It has been shown that the Wnt signaling pathway is dysregulated during viral infections [44], while suppression of apoptosis has also been associated with adenovirus infection [45].

Altogether, our data indicate that HAdV-E4 infection alters the small ncRNA composition of EVs released by infected cells. Like HAdVs of other species, HAdV-E4 encodes two small non-coding RNAs, known as virus associated (VA) RNA_I_ and VA RNA_II_, which are transcribed by RNA Pol III, and are not translated into protein. The VA RNAs are proviral factors that modulate innate host cell defenses and interfere with cellular processes such as nuclear RNA export, protein synthesis and miRNA biogenesis [46, 47]. To investigate whether EVs released by AdV-infected cells carry VA RNAs, the reads obtained from the transcriptomics data were also mapped to the HAdV-E4 genome (GenBank accession #NC_003266.2). Both VA RNA_I_ and VA RNA_II_ were identified in the small RNA cargo of AdV-EVs, with the VA RNA_I_ content being more abundant than that of VA RNA_II_ (Fig. 7A). The observed difference in abundance mirrors the relative VA RNA_I_/VA RNA_II_ abundance reported for HAdV-C-infected cells [46]. The RNA-Seq data for VA RNA_I_/VA RNA_II_ were subsequently validated by RT-qPCR (Fig. 7B).

**Figure 7:**
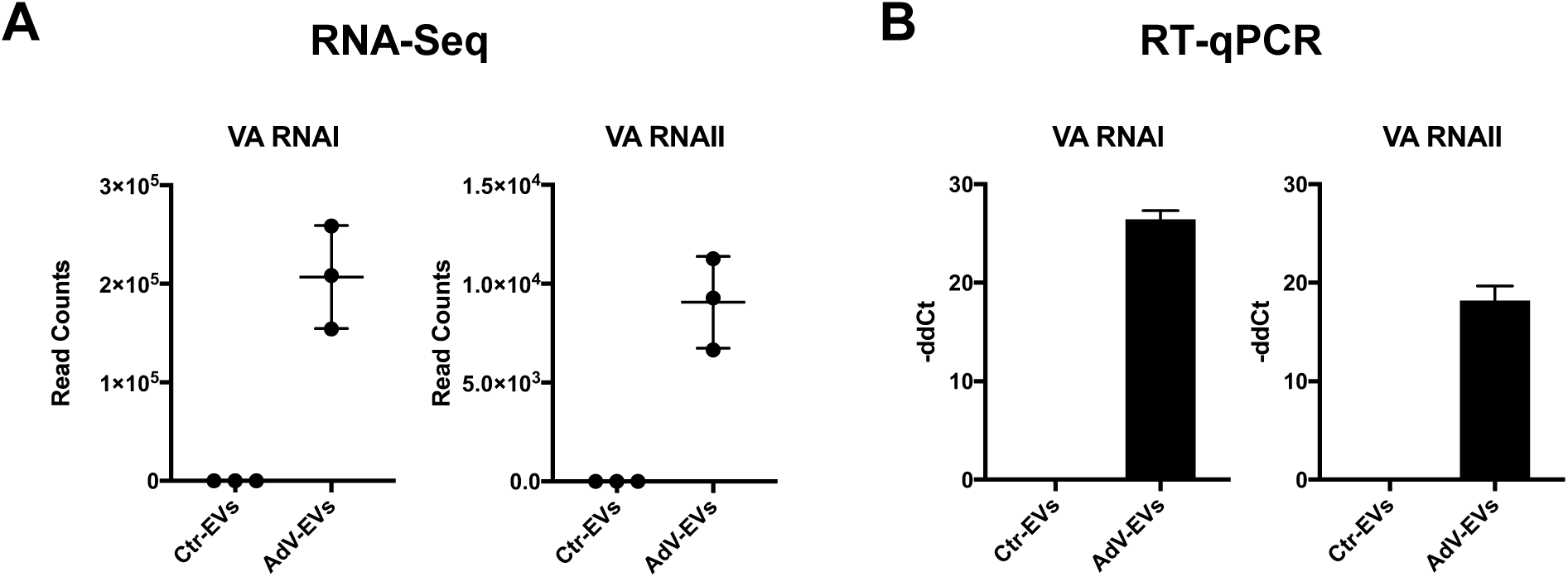
Detection of VA-RNAs in AdV-EVs. **A.** Normalized read counts that aligned to the VA RNAI and VA RNAII sequences of the HAdV-E4 genome. Individual data points of three biological replicates are shown. **B.** RT-qPCR expression data of VA RNA I and VA RNA II in Ctr-EVs and AdV-EVs. Data were normalized on ACTB expression and reported as -ddCt relative to EVs-Ctr. The mean ± SEM from two biological replicates is shown.

### Characterization of EV biological properties

Having shown that AdV-EVs carry viral proteins and viral RNAs, we next tested AdV-EVs for presence of viral genomic DNA by qPCR, and evaluated their infectivity by plaque assay in A549 cells. As shown in Figure 8A, a number of plaques were visible in infected wells indicating that purified EVs from AdV-infected cells carry and successfully transduce replication competent HAdV-E4 genomes to naïve recipient cells. To gain more insights about the localization of EV-associated viral DNA, we subjected AdV-EVs to a series of treatments with DNase I, proteinase K, Triton X-100, and all possible combinations of the these and evaluated their infectivity by plaque assay. We observed a significant reduction in the number of plaques after proteinase K treatment in all treatment combinations. DNase treatment alone was not effective in reducing the number of plaques. Interestingly, treatment with the detergent did not abolish plaque formation, but resulted in the formation of plaques of smaller size (Fig. 8B). The effect of the treatments on the structural integrity of the EVs was also evaluated by TEM. Treatment with DNase I, proteinase K or the combination of the two did not result in any apparent change. In contrast, and as expected, exposure to detergent Triton X-100 lysed all vesicular structures (Fig. 8C). To further evaluate the impact of the various treatments, total DNA was extracted from each treated sample, to determine genome copy concentration by qPCR (Fig. 8D). Reduction of viral DNA load was observed only in samples treated with proteinase K and DNase I, or with Triton X-100, Proteinase K and DNase I. Taking into consideration that a full-length HAdV genome is needed for a productive infection, our data suggest that the viral genome is located inside the EVs, and that proteins present on the surface of the EVs are required for efficient uptake by target cells and subsequent traffic to the nucleus.

**Figure 8:**
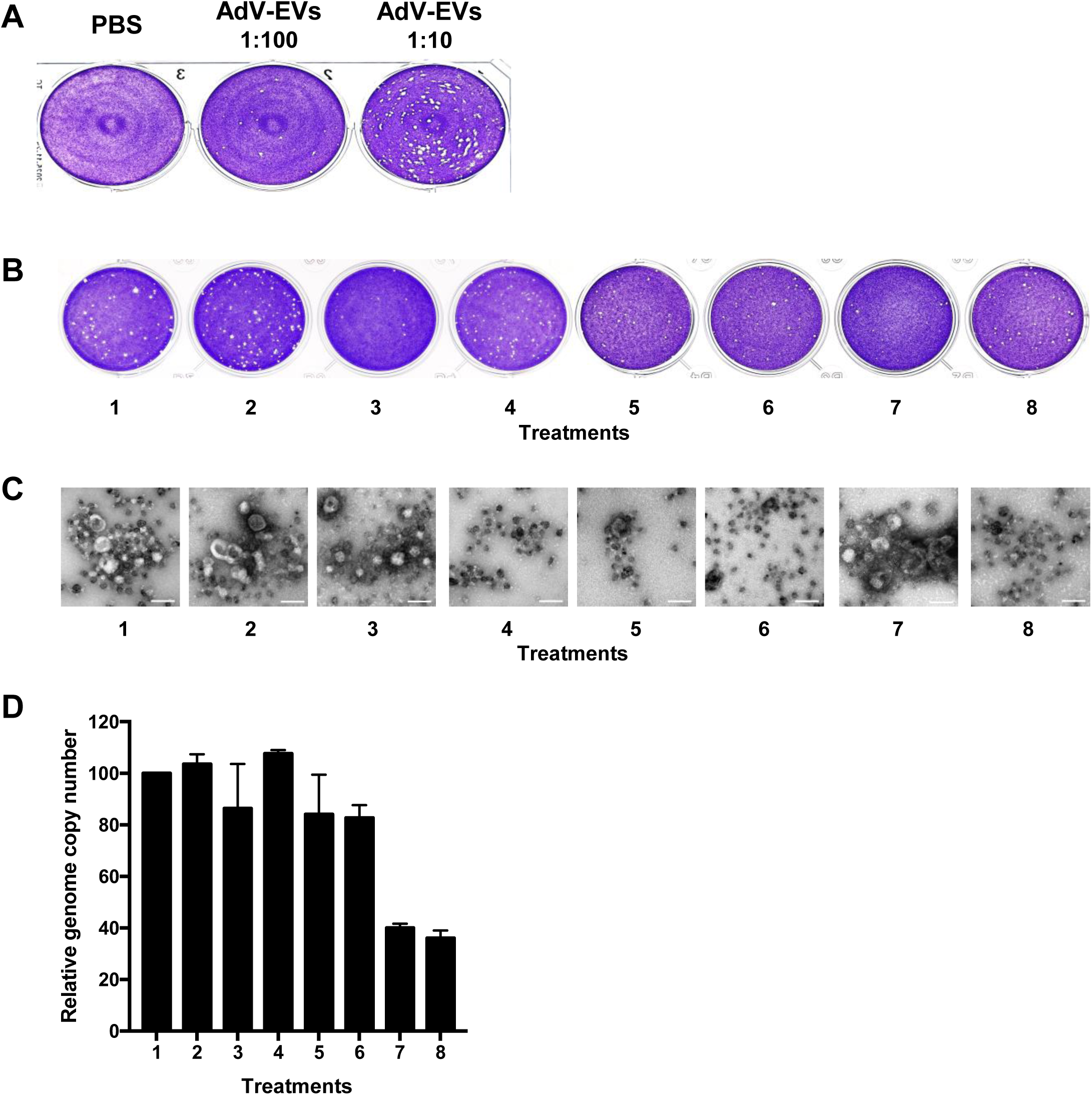
Characterization of EV-associated infectivity. **A.** Plaque assay in A549 cells showing the infectivity of AdV-EVs. **B.** Plaque assay performed with AdV-EV preparations subjected to the following treatments: 1, untreated; 2, DNase I; 3, proteinase K; 4, Triton X-100; 5, Triton X-100/proteinase K; 6, Triton X-100/DNase; 7, proteinase K/DNase I; 8, Triton X-100/proteinase K/DNase I. **C.** TEM images of AdV-EVs treated as in (B). Scale bars = 100 nm. **D.** Genome copy number quantification by qPCR of AdV-EVs treated as indicated above. Bars represent the ratio of genome copy number/mL of treated AdV-EVs over genome copy number/mL of untreated AdV-EVs. Data are shown as mean of two biological replicates ± SEM.

### Neutralization assays

As part of the characterization of the documented infectivity of AdV-EV preps, we carried out plaque reduction neutralization assays using HAdV-E4 virus as a reference for comparison. Interestingly, as shown in Table 6, the rabbit hyperimmune serum S-1001 raised against HAdV-E4 effectively neutralized EV infectivity, completely preventing plaque formation at dilutions ranging from 1:16 to 1:256 and neutralizing ∼70% of EV infectivity at dilution 1:2048. The virus, on the other hand was completely neutralized by serum dilution 1:1024, and ∼70% neutralized by serum dilution 1:8192.

**Table 6:**
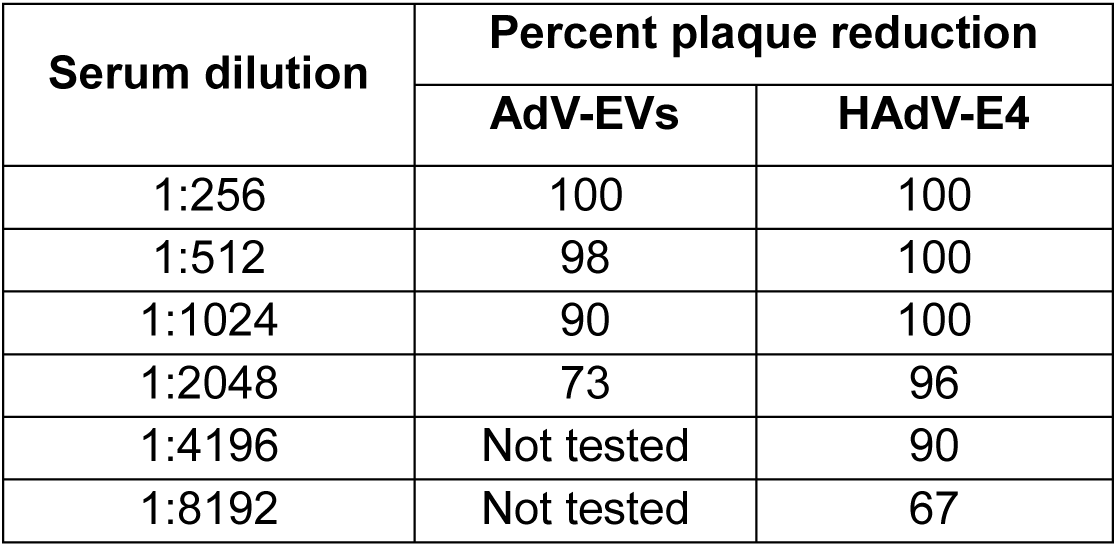
Comparative neutralization of AdV-EV and HAdV-E4 infectivity by rabbit polyclonal anti HAdV-E4 serum.

In an attempt to identify possible EV-associated targets of neutralizing antibodies, protein extracts from Ctr-EVs and AdV-EVs were subjected to immunoprecipitation with the anti-HAdV-E4 rabbit serum and processed for proteomics profiling by mass spectrometry. These experiments confirmed the presence of AdV proteins in AdV-EVs (penton, hexon, pre-hexon-linking protein IIIa and hexon-interlacing protein pIX), and also identified 123 proteins that were present in both Ctr-EVs and AdV-EVs. A GO annotation analysis focused on cellular component (CC) ontology was performed. The results revealed that 23 of the identified proteins are plasma membrane-associated (Table 7) and thus potential targets of neutralizing antibodies.

**Table 7:** Plasma membrane-associated proteins identified by immunoprecipitation of protein extracts of AdV- EVs with rabbit anti-HAdV4 hyperimmune serum and mass spectrometry. GO annotation analysis of cellular component terms.

| Gene ID | Description |
| --- | --- |
| ABCA1 | Phospholipid-transporting ATPase ABCA1 |
| ANXA2 | Annexin A2 |
| APMAP | Adipocyte plasma membrane-associated protein |
| APOA1 | Apolipoprotein A-I |
| ATP2B1 | Plasma membrane calcium-transporting ATPase 1 |
| CD81 | CD81 antigen |
| CTNND1 | Catenin delta-1 |
| DSP | Desmoplakin |
| GNB2 | Guanine nucleotide-binding protein G(I)/G(S)/G(T) subunit beta-2 |
| HP | Haptoglobin |
| IGFBP3 | Insulin-like growth factor-binding protein 3 |
| ITGA3 | Integrin alpha-3 |
| ITGB1 | Integrin beta-1 |
| JUP | Junction plakoglobin |
| KRT77 | Keratin, type II cytoskeletal 1b |
| MYO1B | Unconventional myosin-Ib |
| MYO1C | Unconventional myosin-Ic |
| PKP1 | Plakophilin-1 |
| PLEKHA7 | Pleckstrin homology domain-containing family A member 7 |
| PLG | Plasminogen |
| SLC12A2 | Solute carrier family 12 member 2 |
| SLC3A2 | Amino acid transporter heavy chain SLC3A2 |
| SMOC1 | SPARC-related modular calcium-binding protein 1 |
| STOM | Stomatin |
| TIMP3 | Metalloproteinase inhibitor 3 |

## DISCUSSION

The present study describes the cargo composition of small EVs secreted by HAdV-E4-infected cells at 24h post infection prior to the lytic release of progeny virions, and identifies a potential role for these vesicles in non-lytic cell-to-cell dissemination of infection.

Altogether, the results of the conducted transcriptomics and proteomics analyses indicate that HAdV-E4 infection greatly impacts the content of EVs released by infected cells. While the identified differentially uploaded proteome and small non-coding RNA cargo likely reflect changes in the infected cells occurring in response to infection, they also reveal specific HAdV strategies to hijack EV biogenesis and exploit cargo sorting pathways to upload selected viral-encoded molecules into small EVs increasing the virus’ replicative potential [6]. In support of this hypothesis, we observed dysregulation of proteins involved in the vesicle formation process and in endosomal transport (Fig. 3C), and the presence in AdV-EVs of miRNAs predicted to be involved in such processes (Fig. 6D), although the precise role of let-7f-5p, miR-93-5p, miR-7704 and miR-7706 in adenovirus infection has not been elucidated to the present.

In agreement with the data reported by Saari and colleagues from studies conducted using oncolytic adenovirus vector Ad5/3-D24-GMCSF [17, 48], we identified our AdV-EV population purified using an immunocapture approach as infectious and carrying HAdV-E4 genomic DNA, as well as several virus-encoded proteins. As part of their studies characterizing the role of exosomes in the removal of harmful DNA, Takahashi and colleagues also reported the detection of viral genomic DNA in exosomes secreted from fetal lung fibroblast TIG-3 cells infected with an adenovirus expressing GFP [49]. Consistent with the recent observations of Brachtlova et al. [18], our RNAseq efforts allowed the identification in the cargo of AdV-EVs of the unique viral small non-coding RNAs VA RNA I and VA RNA II.

In the absence of any viral particles detectable in our EV preparations by TEM, we attributed their infectivity to their ability to get uptaken and efficiently deliver capsid-free intact viral genomes to the nucleus of naïve recipient cells. We specifically designed our studies to focus on the EV population released by AdV-infected cells before cell lysis to avoid contamination of our preparations with progeny AdV virions. The integrity of the infected cells at the early time post infection chosen for our harvest (Fig, 1) and the size of our purified EV population rule out the presence of the large cytotoxic vesicles encapsulating mature virions described by Ran and colleagues in their vesicle preparations derived from A549 cells infected with oncolytic Ad5 hTERTp-E1A at 48 h post infection when most cells were detached from the culture dishes [19]. While virions of other virus families have been detected inside small EVs [11, 50], there is currently no evidence that AdV virions can become encapsulated in small (< 200 nm) EVs. However, it has become evident that cells infected by a variety of virus families can secrete EVs containing viral genetic material [51, 52], disclosing a potential role for EVs in non-lytic viral egress and dissemination [11]. Here we showed, for the first time for a replication-competent HAdV type of medical importance, that tetraspanin CD9-, CD63- and CD81-positive small EVs released by infected lung epithelial cells at a pre-lytic early stage during infection carry the viral genome, select structural and non-structural viral proteins and VA RNAs providing a vehicle for protected cell-to-cell dissemination of HAdV infection. Collectively, our data and those of previous studies indicate that AdVs have developed strategies to exploit cellular sorting mechanisms for proteins [53, 54], harmful DNA [49], and miRNAs [55] into small EVs. Given their multiple proviral functions including the inhibition of host innate immune responses through binding of PKR and suppression of the host miRNA biogenesis [47], the presence of VA RNAs inside AdV-EVs appears advantageous to the functional efficiency of these EVs as early vehicles for infection dissemination. This is also the case with DBP, a non-structural protein encoded in the E2A transcriptional unit that plays key roles in viral DNA replication, transcription, and viral gene expression [39], and with non-structural viral shutoff protein L4-100K, which favors translation of viral proteins at the expenses of host proteins [40].

Our characterization of the small non-coding RNA and protein cargo provides a foundation of reference data for future studies investigating the conservation of the identified viral-encoded cargo across members of the seven species comprising HAdV types, and for necessary studies designed to further understand the involvement of EVs during adenovirus infection and their potential contribution to the expansion of cellular/tissue tropism *in vivo*.

Finally, our intriguing findings of neutralization activity of rabbit polyclonal anti HAdV-4 serum against AdV-EV infectivity are consistent with the observations of Saari and colleagues using rabbit polyclonal anti-HAdV-5 antibody on EVs purified from Ad5/3-D24-GMCSF-infected cells [17]. The presence of neutralizing antibodies in both rabbit sera is likely the result of the use of infected cell lysates -instead of purified virions- for the immunization of rabbits. However, these unexpected observations disclose the existence of potential targets for passive immune therapeutic intervention that warrant further investigation.

## ACKNOWLEDGEMENTS

The authors wish to thank Susan Fort at Lovelace for technical assistance with EV preparation and WB analysis; Tamara Howard at the University of New Mexico for sample processing for TEM, and Sam Mackintosh, IDeA National Resource for Quantitative Proteomics for general support and advice with mass spectrometry. This project was supported by Lovelace Biomedical Research Institute intramural support to A.E.K.

## REFERENCES

1. Welsh, J. A.; Goberdhan, D. C. I.; O’Driscoll, L.; Buzas, E. I.; Blenkiron, C.; Bussolati, B.; Cai, H.; Di Vizio, D.; Driedonks, T. A. P.; Erdbrügger, U.; Falcon-Perez, J. M.; Fu, Q.-L.; Hill, A. F.; Lenassi, M.; Lim, S. K.; Mahoney, M. G.; Mohanty, S.; Möller, A.; Nieuwland, R.; Ochiya, T.; Sahoo, S.; Torrecilhas, A. C.; Zheng, L.; Zijlstra, A.; Abuelreich, S.; Bagabas, R.; Bergese, P.; Bridges, E. M.; Brucale, M.; Burger, D.; Carney, R. P.; Cocucci, E.; Crescitelli, R.; Hanser, E.; Harris, A. L.; Haughey, N. J.; Hendrix, A.; Ivanov, A. R.; Jovanovic-Talisman, T.; Kruh-Garcia, N. A.; Ku’ulei-Lyn Faustino, V.; Kyburz, D.; Lässer, C.; Lennon, K. M.; Lötvall, J.; Maddox, A. L.; Martens-Uzunova, E. S.; Mizenko, R. R.; Newman, L. A.; Ridolfi, A.; Rohde, E.; Rojalin, T.; Rowland, A.; Saftics, A.; Sandau, U. S.; Saugstad, J. A.; Shekari, F.; Swift, S.; Ter-Ovanesyan, D.; Tosar, J. P.; Useckaite, Z.; Valle, F.; Varga, Z.; van der Pol, E.; van Herwijnen, M. J. C.; Wauben, M. H. M.; Wehman, A. M.; Williams, S.; Zendrini, A.; Zimmerman, A. J.; Consortium, M.; Théry, C.; Witwer, K. W., Minimal information for studies of extracellular vesicles (MISEV2023): From basic to advanced approaches. Journal of Extracellular Vesicles 2024, 13, (2), e12404.

2. Kim, K. M.; Abdelmohsen, K.; Mustapic, M.; Kapogiannis, D.; Gorospe, M., RNA in extracellular vesicles. Wiley Interdiscip Rev RNA 2017, 8, (4).

3. Jeppesen, D. K.; Fenix, A. M.; Franklin, J. L.; Higginbotham, J. N.; Zhang, Q.; Zimmerman, L. J.; Liebler, D. C.; Ping, J.; Liu, Q.; Evans, R.; Fissell, W. H.; Patton, J. G.; Rome, L. H.; Burnette, D. T.; Coffey, R. J., Reassessment of Exosome Composition. Cell 2019, 177, (2), 428–445 e18.

4. Dixson, A. C.; Dawson, T. R.; Di Vizio, D.; Weaver, A. M., Context-specific regulation of extracellular vesicle biogenesis and cargo selection. Nat Rev Mol Cell Biol 2023, 24, (7), 454–476.

5. Amin, S.; Massoumi, H.; Tewari, D.; Roy, A.; Chaudhuri, M.; Jazayerli, C.; Krishan, A.; Singh, M.; Soleimani, M.; Karaca, E. E.; Mirzaei, A.; Guaiquil, V. H.; Rosenblatt, M. I.; Djalilian, A. R.; Jalilian, E., Cell Type-Specific Extracellular Vesicles and Their Impact on Health and Disease. International Journal of Molecular Sciences 2024, 25, (5), 2730.

6. Mardi, N.; Haiaty, S.; Rahbarghazi, R.; Mobarak, H.; Milani, M.; Zarebkohan, A.; Nouri, M., Exosomal transmission of viruses, a two-edged biological sword. Cell Commun Signal 2023, 21, (1), 19.

7. Feng, Z.; Hensley, L.; McKnight, K. L.; Hu, F.; Madden, V.; Ping, L.; Jeong, S.-H.; Walker, C.; Lanford, R. E.; Lemon, S. M., A pathogenic picornavirus acquires an envelope by hijacking cellular membranes. Nature 2013, 496, (7445), 367–371.

8. Huang, H.-I.; Lin, J.-Y.; Chiang, H.-C.; Huang, P.-N.; Lin, Q.-D.; Shih, S.-R., Exosomes Facilitate Transmission of Enterovirus A71 From Human Intestinal Epithelial Cells. The Journal of Infectious Diseases 2020, 222, (3), 456–469.

9. Robinson, S. M.; Tsueng, G.; Sin, J.; Mangale, V.; Rahawi, S.; McIntyre, L. L.; Williams, W.; Kha, N.; Cruz, C.; Hancock, B. M.; Nguyen, D. P.; Sayen, M. R.; Hilton, B. J.; Doran, K. S.; Segall, A. M.; Wolkowicz, R.; Cornell, C. T.; Whitton, J. L.; Gottlieb, R. A.; Feuer, R., Coxsackievirus B exits the host cell in shed microvesicles displaying autophagosomal markers. PLoS Pathog 2014, 10, (4), e1004045.

10. Iša, P.; Pérez-Delgado, A.; Quevedo, I. R.; López, S.; Arias, C. F., Rotaviruses Associate with Distinct Types of Extracellular Vesicles. Viruses 2020, 12, (7).

11. Yang, J. E.; Rossignol, E. D.; Chang, D.; Zaia, J.; Forrester, I.; Raja, K.; Winbigler, H.; Nicastro, D.; Jackson, W. T.; Bullitt, E., Complexity and ultrastructure of infectious extracellular vesicles from cells infected by non-enveloped virus. Scientific Reports 2020, 10, (1), 7939.

12. Lynch, J. P., 3rd; Kajon, A. E., Adenovirus: Epidemiology, Global Spread of Novel Types, and Approach to Treatment. Semin Respir Crit Care Med 2021, 42, (6), 800–821.

13. Kajon, A. E., Adenovirus infections: new insights for the clinical laboratory. J Clin Microbiol 2024, 62, (9), e0083622.

14. Lion, T., Adenovirus persistence, reactivation, and clinical management. FEBS Lett 2019, 593, (24), 3571–3582.

15. MacNeil, K. M.; Dodge, M. J.; Evans, A. M.; Tessier, T. M.; Weinberg, J. B.; Mymryk, J. S., Adenoviruses in medicine: innocuous pathogen, predator, or partner. Trends Mol Med 2023, 29, (1), 4–19.

16. Ipinmoroti, A. O.; Crenshaw, B. J.; Pandit, R.; Kumar, S.; Sims, B.; Matthews, Q. L., Human Adenovirus Serotype 3 Infection Modulates the Biogenesis and Composition of Lung Cell-Derived Extracellular Vesicles. J Immunol Res 2021, 2021, 2958394.

17. Saari, H.; Turunen, T.; Lõhmus, A.; Turunen, M.; Jalasvuori, M.; Butcher, S. J.; Ylä-Herttuala, S.; Viitala, T.; Cerullo, V.; Siljander, P. R. M.; Yliperttula, M., Extracellular vesicles provide a capsid-free vector for oncolytic adenoviral DNA delivery. J Extracell Vesicles 2020, 9, (1), 1747206.

18. Brachtlova, T.; Li, J.; van der Meulen-Muileman, I. H.; Sluiter, F.; von Meijenfeldt, W.; Witte, I.; Massaar, S.; van den Oever, R.; de Vrij, J.; van Beusechem, V. W., Quantitative Virus-Associated RNA Detection to Monitor Oncolytic Adenovirus Replication. Int J Mol Sci 2024, 25, (12).

19. Ran, L.; Tan, X.; Li, Y.; Zhang, H.; Ma, R.; Ji, T.; Dong, W.; Tong, T.; Liu, Y.; Chen, D.; Yin, X.; Liang, X.; Tang, K.; Ma, J.; Zhang, Y.; Cao, X.; Hu, Z.; Qin, X.; Huang, B., Delivery of oncolytic adenovirus into the nucleus of tumorigenic cells by tumor microparticles for virotherapy. Biomaterials 2016, 89, 56–66.

20. Garofalo, M.; Saari, H.; Somersalo, P.; Crescenti, D.; Kuryk, L.; Aksela, L.; Capasso, C.; Madetoja, M.; Koskinen, K.; Oksanen, T.; Makitie, A.; Jalasvuori, M.; Cerullo, V.; Ciana, P.; Yliperttula, M., Antitumor effect of oncolytic virus and paclitaxel encapsulated in extracellular vesicles for lung cancer treatment. J Control Release 2018, 283, 223–234.

21. Garofalo, M.; Villa, A.; Rizzi, N.; Kuryk, L.; Rinner, B.; Cerullo, V.; Yliperttula, M.; Mazzaferro, V.; Ciana, P., Extracellular vesicles enhance the targeted delivery of immunogenic oncolytic adenovirus and paclitaxel in immunocompetent mice. J Control Release 2019, 294, 165–175.

22. Bair, C. R.; Zhang, W.; Gonzalez, G.; Kamali, A.; Stylos, D.; Blanco, J. C. G.; Kajon, A. E., Human Adenovirus Type 4 Comprises Two Major Phylogroups with Distinct Replicative Fitness and Virulence Phenotypes. Journal of Virology 2022, 96, (5), e01090–21.

23. Hughes, C. S.; Moggridge, S.; Müller, T.; Sorensen, P. H.; Morin, G. B.; Krijgsveld, J., Single-pot, solid-phase-enhanced sample preparation for proteomics experiments. Nature Protocols 2019, 14, (1), 68–85.

24. Graw, S.; Tang, J.; Zafar, M. K.; Byrd, A. K.; Bolden, C.; Peterson, E. C.; Byrum, S. D., proteiNorm - A User-Friendly Tool for Normalization and Analysis of TMT and Label-Free Protein Quantification. ACS Omega 2020, 5, (40), 25625–25633.

25. Huber, W.; von Heydebreck, A.; Sultmann, H.; Poustka, A.; Vingron, M., Variance stabilization applied to microarray data calibration and to the quantification of differential expression. Bioinformatics 2002, 18 Suppl 1, S96–104.

26. Thurman, T. J.; Washam, C. L.; Alkam, D.; Bird, J. T.; Gies, A.; Dhusia, K.; Robeson, M. S.; Byrum, S. D., proteoDA: a package for quantitative proteomics. Journal of Open Source Software 2023, 8, (85), 5184.

27. Ritchie, M. E.; Phipson, B.; Wu, D.; Hu, Y.; Law, C. W.; Shi, W.; Smyth, G. K., limma powers differential expression analyses for RNA-sequencing and microarray studies. Nucleic Acids Res 2015, 43, (7), e47.

28. Keerthikumar, S.; Chisanga, D.; Ariyaratne, D.; Al Saffar, H.; Anand, S.; Zhao, K.; Samuel, M.; Pathan, M.; Jois, M.; Chilamkurti, N.; Gangoda, L.; Mathivanan, S., ExoCarta: A Web-Based Compendium of Exosomal Cargo. Journal of Molecular Biology 2016, 428, (4), 688–692.

29. Chitti, S. V.; Gummadi, S.; Kang, T.; Shahi, S.; Marzan, A. L.; Nedeva, C.; Sanwlani, R.; Bramich, K.; Stewart, S.; Petrovska, M.; Sen, B.; Ozkan, A.; Akinfenwa, M.; Fonseka, P.; Mathivanan, S., Vesiclepedia 2024: an extracellular vesicles and extracellular particles repository. Nucleic Acids Research 2023, 52, (D1), D1694–D1698.

30. Kang, J.; Tang, Q.; He, J.; Li, L.; Yang, N.; Yu, S.; Wang, M.; Zhang, Y.; Lin, J.; Cui, T.; Hu, Y.; Tan, P.; Cheng, J.; Zheng, H.; Wang, D.; Su, X.; Chen, W.; Huang, Y., RNAInter v4.0: RNA interactome repository with redefined confidence scoring system and improved accessibility. Nucleic Acids Res 2022, 50, (D1), D326–D332.

31. Heim, A.; Ebnet, C.; Harste, G.; Pring-Åkerblom, P., Rapid and quantitative detection of human adenovirus DNA by real-time PCR. Journal of Medical Virology 2003, 70, (2), 228–239.

32. Rubinstein, E.; Thery, C.; Zimmermann, P., Tetraspanins affect membrane structures and the trafficking of molecular partners: what impact on extracellular vesicles? Biochem Soc Trans 2025, 0, (0), 371–82.

33. Szklarczyk, D.; Kirsch, R.; Koutrouli, M.; Nastou, K.; Mehryary, F.; Hachilif, R.; Gable, A. L.; Fang, T.; Doncheva, N. T.; Pyysalo, S.; Bork, P.; Jensen, L. J.; von Mering, C., The STRING database in 2023: protein-protein association networks and functional enrichment analyses for any sequenced genome of interest. Nucleic Acids Res 2023, 51, (D1), D638–d646.

34. Kawabe, T.; Matsushima, M.; Hashimoto, N.; Imaizumi, K.; Hasegawa, Y., CD40/CD40 ligand interactions in immune responses and pulmonary immunity. Nagoya J Med Sci 2011, 73, (3-4), 69–78.

35. O’Shea, C. C.; Choi, S.; McCormick, F.; Stokoe, D., Adenovirus overrides cellular checkpoints for protein translation. Cell Cycle 2005, 4, (7), 883–8.

36. Jung, B. K.; Kim, Y. J.; Hong, J.; Chang, H. G.; Yoon, A. R.; Yun, C. O., ErbB3-Targeting Oncolytic Adenovirus Causes Potent Tumor Suppression by Induction of Apoptosis in Cancer Cells. Int J Mol Sci 2022, 23, (13).

37. Albarnaz, J. D.; Weekes, M. P., Proteomic analysis of antiviral innate immunity. Curr Opin Virol 2023, 58, 101291.

38. Mellors, J.; Tipton, T.; Longet, S.; Carroll, M., Viral Evasion of the Complement System and Its Importance for Vaccines and Therapeutics. Frontiers in Immunology 2020, 11.

39. Bertzbach, L. D.; Seddar, L.; von Stromberg, K.; Ip, W. H.; Dobner, T.; Hidalgo, P., The adenovirus DNA-binding protein DBP. J Virol 2024, 98, (2), e0188523.

40. Weitzman, M. D.; Ornelles, D. A., Inactivating intracellular antiviral responses during adenovirus infection. Oncogene 2005, 24, (52), 7686–7696.

41. Murray, J. L.; Sheng, J.; Rubin, D. H., A role for H/ACA and C/D small nucleolar RNAs in viral replication. Mol Biotechnol 2014, 56, (5), 429–37.

42. Price, A. M.; Hayer, K. E.; McIntyre, A. B. R.; Gokhale, N. S.; Abebe, J. S.; Della Fera, A. N.; Mason, C. E.; Horner, S. M.; Wilson, A. C.; Depledge, D. P.; Weitzman, M. D., Direct RNA sequencing reveals m(6)A modifications on adenovirus RNA are necessary for efficient splicing. Nat Commun 2020, 11, (1), 6016.

43. Hajikhezri, Z.; Kaira, Y.; Schubert, E.; Darweesh, M.; Svensson, C.; Akusjarvi, G.; Punga, T., Fragile X-Related Protein FXR1 Controls Human Adenovirus Capsid mRNA Metabolism. J Virol 2023, 97, (2), e0153922.

44. van Zuylen, W. J.; Rawlinson, W. D.; Ford, C. E., The Wnt pathway: a key network in cell signalling dysregulated by viruses. Reviews in Medical Virology 2016, 26, (5), 340–355.

45. Zhao, H.; Granberg, F.; Pettersson, U., How adenovirus strives to control cellular gene expression. Virology 2007, 363, (2), 357–375.

46. Punga, T.; Darweesh, M.; Akusjärvi, G., Synthesis, Structure, and Function of Human Adenovirus Small Non-Coding RNAs. Viruses 2020, 12, (10), 1182.

47. Vachon, V. K.; Conn, G. L., Adenovirus VA RNA: An essential pro-viral non-coding RNA. Virus Res 2016, 212, 39–52.

48. Koski, A.; Kangasniemi, L.; Escutenaire, S.; Pesonen, S.; Cerullo, V.; Diaconu, I.; Nokisalmi, P.; Raki, M.; Rajecki, M.; Guse, K.; Ranki, T.; Oksanen, M.; Holm, S. L.; Haavisto, E.; Karioja-Kallio, A.; Laasonen, L.; Partanen, K.; Ugolini, M.; Helminen, A.; Karli, E.; Hannuksela, P.; Pesonen, S.; Joensuu, T.; Kanerva, A.; Hemminki, A., Treatment of cancer patients with a serotype 5/3 chimeric oncolytic adenovirus expressing GMCSF. Mol Ther 2010, 18, (10), 1874–84.

49. Takahashi, A.; Okada, R.; Nagao, K.; Kawamata, Y.; Hanyu, A.; Yoshimoto, S.; Takasugi, M.; Watanabe, S.; Kanemaki, M. T.; Obuse, C.; Hara, E., Exosomes maintain cellular homeostasis by excreting harmful DNA from cells. Nat Commun 2017, 8, 15287.

50. Wu, Q.; Glitscher, M.; Tonnemacher, S.; Schollmeier, A.; Raupach, J.; Zahn, T.; Eberle, R.; Krijnse-Locker, J.; Basic, M.; Hildt, E., Presence of Intact Hepatitis B Virions in Exosomes. Cell Mol Gastroenterol Hepatol 2023, 15, (1), 237–259.

51. Chatterjee, S.; Kordbacheh, R.; Sin, J., Extracellular Vesicles: A Novel Mode of Viral Propagation Exploited by Enveloped and Non-Enveloped Viruses. Microorganisms 2024, 12, (2).

52. Kumar, A.; Kodidela, S.; Tadrous, E.; Cory, T. J.; Walker, C. M.; Smith, A. M.; Mukherjee, A.; Kumar, S., Extracellular Vesicles in Viral Replication and Pathogenesis and Their Potential Role in Therapeutic Intervention. Viruses 2020, 12, (8).

53. Chen, Y.; Zhao, Y.; Yin, Y.; Jia, X.; Mao, L., Mechanism of cargo sorting into small extracellular vesicles. Bioengineered 2021, 12, (1), 8186–8201.

54. Waury, K.; Gogishvili, D.; Nieuwland, R.; Chatterjee, M.; Teunissen, C. E.; Abeln, S., Proteome encoded determinants of protein sorting into extracellular vesicles. J Extracell Biol 2024, 3, (1), e120.

55. Groot, M.; Lee, H., Sorting Mechanisms for MicroRNAs into Extracellular Vesicles and Their Associated Diseases. Cells 2020, 9, (4).

